# Simultaneous representation of sensory and mnemonic information in human visual cortex

**DOI:** 10.1101/339200

**Authors:** R. L. Rademaker, C. Chunharas, J. T. Serences

**Affiliations:** Psychology Department, University of California San Diego, La Jolla, California, USA; Donders Institute for Brain, Cognition and Behavior, Radboud University, Nijmegen, the Netherlands; Neurosciences Graduate Program, University of California San Diego, La Jolla, California, USA; Kavli Institute for Brain and Mind, University of California, San Diego, La Jolla, CA 92093

## Abstract

Traversing sensory environments requires keeping relevant information in mind while simultaneously processing new inputs. Visual information is kept in working memory via feature selective responses in early visual cortex, but recent work had suggested that new sensory inputs wipe out this information. Here we show region-wide multiplexing abilities in classic sensory areas, with population-level response patterns in visual cortex representing the contents of working memory concurrently with new sensory inputs.

---

When trying to attain behavioral goals, the ability to flexibly juggle thoughts is key. Visual Working Memory (VWM) provides the mental workspace to keep visual information online, allowing this information to guide visual search or to be recalled at a future moment in time. Neuroimaging studies have firmly established that VWM contents can be decoded from occipital visual areas, including primary visual cortex V1 (Harrison & Tong, 2009; Serences, Ester, Vogel, & Awh, 2009; Riggal & Postle, 2012; Christophel, Hebart, & Haynes, 2012). The quality of this information can predict behavioral performance Ester, Anderson, Serences, & Awh, 2013; Bettencourt & Xu, 2016), suggesting that early sensory areas are involved in the representation of visual memories.

That said, previous studies typically relied on a traditional delayed-match-to-sample (DMTS) task in which a sample memory stimulus is encoded and remembered across a blank delay interval before a test stimulus appears for comparison. However, in everyday perception, VWM maintenance needs to be robust to the continuous influx of new visual inputs that come with each exploratory saccade or change in the environment. Thus, a delay period devoid of other visual inputs is quite divorced from typical visual experience. Based on this mismatch between experimental and real-world scenarios, some have argued that recruiting early sensory areas to store relevant VWM information would be counterproductive in everyday life, as new sensory inputs would destructively interfere with concurrent mnemonic representations (Bettencourt & Xu, 2016; Mendoza-Halliday, Torres, & Martinez-Trujillo, 2014; Stokes, 2015).

Based on this logic, one recent study (Bettencourt & Xu, 2016) employed a DMTS task with task-irrelevant pictures of faces and gazebos sometimes presented during the delay period. The authors used functional magnetic resonance imaging (fMRI) and found that activation patterns in early visual cortex represented the contents of VWM during blank delays, but not when delays were filled with predictable distractors (i.e. the task-irrelevant pictures). The authors concluded that representations initially encoded in early visual cortex were recoded in a more durable format in parietal cortex to insulate mnemonic representations from interference by new sensory input. Furthermore, the authors argued that the disengagement of primary sensory regions was strategic as it occurred only when participants expected the task-irrelevant pictures. This study challenged the role of early visual cortex during VWM, and would suggest that previous findings of sensory recruitment can be an artifact of overly artificial tasks.

Importantly, the framework proposed by Bettencourt and Xu (2016) implies a fundamental limitation of cortical information processing: a sensory area such as primary visual cortex cannot simultaneously represent top-down biasing signals associated with internal cognitive operations like VWM and bottom-up sensory inputs evoked by newly encountered stimuli in the environment. However, there are at least two reasons to question the strongest version of this stance. First, bottom-up input from the lateral geniculate nucleus primarily projects to layer 4 of primary visual cortex, whereas top-down input arrives primarily in superficial layers (Nassi & Callaway, 2009; Van Kerkoerle, Self, & Roelfsema, 2017). This anatomical segregation of bottom-up and top-down inputs could theoretically support the co-existence of multiple simultaneous representations, a concept we term ‘region-wide multiplexing’ (after the more common usage of ‘multiplexing’ to refer to flexible coding in single neurons). Second, from a functional point of view, success on a DMTS task relies on comparing an internally stored representation to a new sensory input. For this, a ‘local comparison circuit’, able to jointly represent remembered and perceived items in the same local circuit, could be ideal.

To evaluate the ability of early visual areas to act as a multiplexing comparison circuit during VWM, participants remembered a randomly oriented visual grating for 13 seconds while looking at either a blank screen or a sequence of visual distractors (**Fig. 1a**). Distractors were either a Fourier filtered white noise stimulus, or another orientated grating with random angular offset relative to the memory orientation. Distractors were predictable, but irrelevant to the task, and their presence had no observable impact on behavior during fMRI scanning (**Supplementary Fig. 1**). Note however that, in a separate behavioral experiment using a similar paradigm with many more trials, the orientation of a distractor systematically biased recall of a remembered orientation (**Supplementary Fig. 2**). As expected, distractors effectively drove a sustained and highly robust increase in the overall univariate response amplitude in V1 and other early visual areas (**Supplementary Fig. 3**).

**Figure 1.**
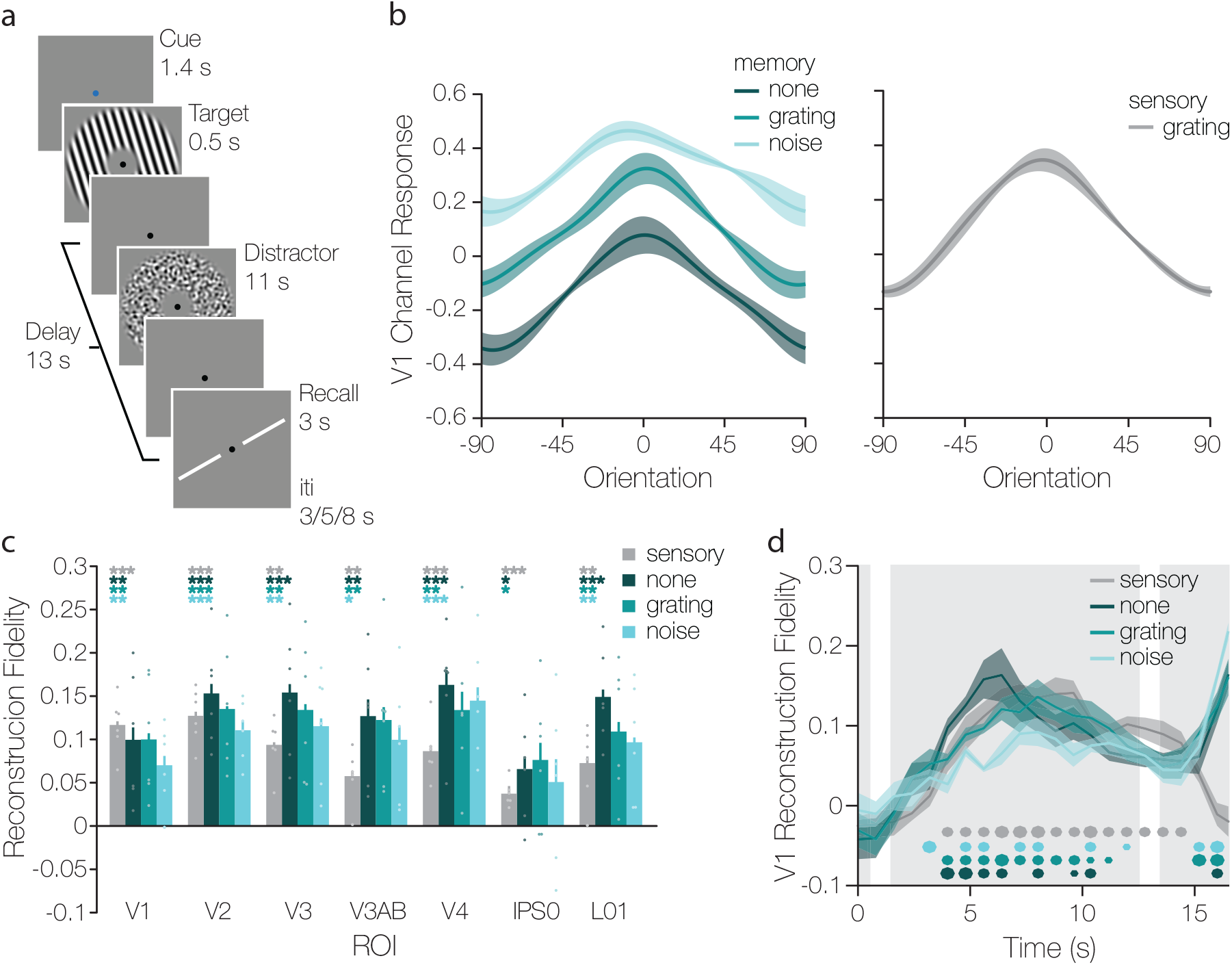
Reconstructions of remembered orientations and oriented distractors during the memory delay period (**a**) Participants were validly cued which distractor to expect during the delay (here, the blue fixation cued a noise distractor). Next a target orientation was shown, and remembered throughout a 13-second delay. Participants viewed either a grey screen, or an 11-second counter phase flickering distractor during the middle portion of the delay. Distractors could be an oriented grating (depicted) or Fourier filtered noise. After the memory delay participants had 3 seconds to rotate a recall probe to match the remembered orientation as precisely as possible. (**b**) Reconstructions of the remembered orientation during the three different distractor conditions (left), and of the physically-present sensory orientation on trials with a grating distractor (right). Reconstructions were based on the average activation patterns from 5.6–13.6 seconds after stimulus onset. (**c**) The degree to which memory and sensory stimuli were represented during the delay was quantified by projecting the channel response at each degree onto a vector centered on the remembered orientation (i.e. zero). The mean of these vectors represents the amount of information for the remembered orientation (see Online Methods). One, two, or three asterisks indicate significance levels of p < 0.05, p < 0.01, and p < 0.001, respectively (based on randomization tests comparing the fidelity in each condition and ROI to 0, see Online Methods). (**d**) Fidelity of timepoint-by-timepoint reconstructions in V1, with time 0 representing target onset. The three gray panels represent the target, distractor, and recall epochs of the working memory trial. Small, medium, and large dots at the bottom indicate significance at each time point at p ≤ 0.05, p ≤ 0.01, and p ≤ 0.001, respectively. Error bars/ areas all represent ± 1 within-subject SEM.

Next, using a multivariate ‘inverted encoding model’ Brouwer & Heeger, 2009; Sprague, Saproo, & Serences, 2015) (trained on independent data, **Supplementary Fig. 4**) we could reconstruct the orientation in memory from delay period activity patterns in primary visual cortex (**Fig. 1b**, left panel), and all other areas along the visual hierarchy (**Supplementary Fig. 5 and 6**), irrespective of whether a distractor was present during the delay. The baseline offset observed between distractor conditions (vertical shift up/down the y-axis in **Fig. 1b**) largely reflects the univariate effect of distractor presence during the delay interval, with higher baselines during trials with distractors (see **Supplementary Fig. 3**). As a measure of tuning fidelity, we projected the vector length of the channel response at each degree in orientation space onto the remembered orientation (i.e. zero in **Fig. 1b**), and calculated the mean. Projected vector means were significantly above zero, indicating that there was information about the remembered orientation during all distractor conditions, and in all regions of interest (ROIs) except IPS0 (**Fig. 1C**, teal bars). Fidelity differed between distractor conditions (F_(2,10)_ = 8.832, p = 0.001), being lower when a noise distractor was present compared to no distractor or a grating distractor (all p < = 0.01; **Supplementary Table 1**). Fidelity also differed between ROIs (F_(6,30)_ = 4.836, p = 0.01), with V1 having lower fidelity than V2, V3, V4, and L01 (all p’s < 0.03; **Supplementary Table 2**), and IPS0 having lower fidelity than V2 and LO1 (both p < 0.001).

Importantly, we were also able to simultaneously reconstruct the orientation of the distractor that was physically present on the screen during one-third of trials (**Fig. 1b**, right panel; **Fig. 1c**, grey bars). This demonstrates that, contrary to simple feedforward models that posit V1 as a passive filter, early visual cortex can concurrently represent incoming sensory information alongside mnemonic information that is no longer tethered to external inputs. While the fidelity of concurrently remembered and sensed orientations was roughly equivalent in V1–V2 (**Fig. 1c**), the fidelity of the sensory grating dropped against the fidelity of the memory grating when ascending the visual hierarchy (interaction F_(6,30)_ = 3.2, p = 0.007). This finding captures the top-down nature of the mnemonic signals, as top-down signals are thought to have more traction than bottom-up sensory inputs in higher-level regions. Importantly, timepoint-by-timepoint reconstructions reveal that remembered and perceived representations evolve concurrently over time in primary visual cortex (**Fig. 1d**) and all along the visual hierarchy (**Supplementary Fig. 7 and 8**), indicating that these representations coexist throughout most of the delay period. The notion of a comparison circuit at the level of early sensory cortex is further supported by a boost in representational quality when target and distractor orientations are similar, compared to dissimilar (**Supplementary Fig. 9**). This finding is in line with behavioral and neuroimaging demonstrations showing higher fidelity memory recall for similar targets and distractors (Rademaker, Bloem, De Weerd, & Sack, 2015; Lorenc, Sreenivasan, Nee, Vandenbroucke, & D’Esposito, 2018; **Supplementary Fig. 2**).

The coexistence of feature-selective top-down inputs that carry information about remembered items, *combined* with local information about current sensory inputs, could provide a powerful local mechanism for comparing memory contents to the sensory environment (Bisley, Zaksas, Droll, & Pasternak, 2004; Zaksas & Pasternak, 2006). For example, if neurons tuned to the features of the sample stimulus were selectively activated during the delay, the output of a local comparison circuit would be relatively high when a matching test stimulus was encountered that selectively excited the same set of neurons. On the other hand, a mismatch test stimulus would drive a new set of neurons, which may lead to a lower overall response due to inhibitory competition with the already active neurons tuned to the sample feature (Serences, 2016; Gayet, et al., 2017). The same logic also applies if the top-down modulations supporting VWM do not lead to sustained patterns of spiking in sensory cortices, and instead only influence sub-threshold potentials (Mendoza-Halliday, Torres, & Martinez-Trujillo, 2014; Merrikhi et al., 2017). Differences in the local comparison circuit output would still be expected due to interactions between the top-down feature-selective bias and the sensory response evoked by the test stimulus. Moreover, content-specific patterns of sub-threshold membrane potentials could persevere through bouts of local spiking driven by sensory inputs during a delay, and thus protect mnemonic contents from distraction (Serences, 2016).

## Acknowledgements

This work was supported by NEI R01-EY025872 and a James S McDonnell Foundation Scholar Award to JTS, and by the European Union’s Horizon 2020 research and innovation program under the Marie Sklodowska-Curie Grant Agreement No 743941 to RLR. We thank Aaron Jacobson at the UCSD center for functional Magnetic Resonance Imaging (CFMRI) for assistance with multi-band imaging protocols. We also thank Ruben van Bergen for assistance setting up an FSL/FreeSurfer retinotopy pipeline, Ahana Chakraborty for collecting the behavioral data, Vy Vo for discussions on statistical analyses, and Stephanie Nelli for feedback on the manuscript.

## Online Methods

### Participants

Six volunteers (5 female) between the ages of 21 and 32 years participated in this experiment at the University of California San Diego. The local Institutional Review Board approved the experiment. All participants provided written informed consent, had normal or corrected-to-normal vision, and received monetary reimbursement ($20 an hour) for their time.

### Stimuli

All stimuli were projected on a 120 x 90 cm screen placed at the foot-end of the scanner and viewed through a front-facing mirror from ~370 cm in an otherwise darkened room. Stimuli were generated under OS X using MATLAB 2013a and the Psychophysics toolbox^21,22^. The luminance output from the projector was linearized, and all stimuli were presented against a 62.82 cd/m^2^ uniform grey background. Stimuli consisted of donut-shaped (1.5° and 7° inner and outer radius, respectively) or circular (1.5° radius) sinusoidal gratings with smoothed edges (1° kernel; sd = 0.5°). In addition to grating stimuli, donut-shaped distractors could also be Fourier filtered noise stimuli, consisting of white noise filtered to include only spatial frequencies between 1 and 4 c/°. The contrast of the distractors was 50% Michelson, while memory targets were presented at full contrast. Stimuli presented during the memory and mapping tasks were counter phase flickering at 4 Hz and 5 Hz, respectively. All sinusoidal gratings had a spatial frequency of 2 c/° and an orientation that was chosen at random from one of six orientation bins. Each orientation bin contained 30 orientations, in integers, and orientations were drawn from each bin equally often. This was done to ensure a relatively uniform sampling of orientation space. A 0.2° central dot aided fixation throughout (black during the memory task, and mean grey with a 0.1° radius magenta dot on top during the mapping task). The recall probe consisted of two 0.035° wide white line segments, with each 5.5° long segment presented at the same distance from fixation as the donut-shaped stimuli.

### Procedure

In the memory task (**Fig. 1a**), each trial started with a 1.4 second change in the color of the central fixation dot, indicating the distractor condition during the delay (e.g. no distractor, noise distractor, grating distractor). Cues could be blue, green, or red, and cue luminances were equated outside the scanner using heterochromatic flicker fusion. Cue color and distractor condition pairings were randomized across participants. A memory target was shown for 500 ms and participants remembered its orientation over a 13 second delay. A noise or grating distractor was presented for 11 seconds during the middle portion of the delay (starting 1s after the offset of the memory target), or the screen remained grey throughout. Importantly, on trials with a grating distractor, target and distractor orientations were selected independently of one another. After the delay, participants used four buttons to rotate a recall probe around fixation, matching the orientation in memory as precisely as possible. The left two buttons rotated the line counter clockwise, while the right two buttons rotated it clockwise. Using the outer- or inner-most buttons would result in faster or slower rotation of the recall probe, respectively. Participants had 3 seconds to respond before being presented with the next memory target 3, 5, or 8 seconds later. Each run consisted of 12 memory trials, and lasted 4 minutes and 40.8 seconds. Distractor type (grating, noise, or none) and the orientation bin from which the target or distractor grating orientations were drawn (one of six), were fully counterbalanced across 9 consecutive runs of the memory task.

During the independent mapping/localizer task, participants viewed 9-second blocks of donut-shaped or circle-shaped grating stimuli. Per run, both stimulus types were alternated 20 times, with 4 fixation blocks interspersed. To ensure participants attended the physical location of the stimuli, they performed a detection task: Grating contrast was probabilistically dimmed twice every 9 seconds, from 100% to 80% for 200 ms. Because the contrast change was probabilistic, some stimulus blocks saw no change, while others had >2 changes. Note that the donut-shaped stimuli in the mapping task occupied the same physical location as the donut-shaped stimuli during the memory task. A mapping run lasted 7 minutes.

Both tasks were practiced outside the scanner before the first scanning session. For the memory task, practice lasted until participants were comfortable using the buttons to recall orientation within the temporally restricted response window, and until their mean absolute error on a run was <10°. For all participants, the data presented here were acquired during 3 separate scanning sessions. During each session, participants completed 8–10 runs of the memory task (18 total runs across days), and 4–6 runs of the mapping task (15–17 total runs across days).

### Magnetic Resonance Imaging

All scans were performed on a General Electric (GE) Discovery MR750 3.0T scanner located at the University of California, San Diego (UCSD), Keck Center for Functional Magnetic Resonance Imaging (CFMRI). High resolution (1 mm^3^ isotropic) anatomical images were acquired during a retinotopic mapping session, using an Invivo 8-channel head coil. Functional echo-planar imaging (EPI) data for the current experiment were acquired using a Nova Medical 32-channel head coil (NMSC075-32-3GE-MR750) and the Stanford Simultaneous Multi-Slice (SMS) EPI sequence (MUX EPI), utilizing 9 axial slices per band and a multiband factor of 8 (total slices = 72; 2 mm^3^ isotropic; 0 mm gap; matrix = 104 x 104; FOV = 20.8 cm; TR/TE = 800/35 ms, flip angle = 52°; inplane acceleration = 1). At sequence onset, the initial 16 TR’s served as reference images critical to the transformation from k-space to image space. Un-aliasing and image reconstruction procedures were performed on local servers using CNI based reconstruction code. Forward and reverse phase-encoding directions were utilized during the acquisition of two short (17 s) “topup” datasets. From these images, susceptibility-induced off-resonance fields were estimated^23^ and used to correct signal distortion inherent in EPI sequences using FSL topup^24,25^.

### Preprocessing

All imaging data were preprocessed using software tools developed and distributed by FreeSurfer and FSL (free to download at https://surfer.nmr.mgh.harvard.edu and http://www.fmrib.ox.ac.uk/fsl). Cortical surface gray-white matter volumetric segmentation of the high resolution anatomical image was performed using the “recon-all” utility in the FreeSurfer analysis suite^26^. Segmented T1 data were used to define Regions of Interest (ROIs) for use in subsequent analyses. The first volume of every functional run was then coregistered to this common anatomical image. Transformation matrices were generated using FreeSurfer’s manual and boundary based registration tools^27^. These matrices were then used to transform each 4D functional volume using FSL FLIRT^28,29^, such that all cross-session data from a single participant was in the same space. Next, motion correction was performed using the FSL tool MCFLIRT^29^ without spatial smoothing, a final sinc interpolation stage, and 12 degrees of freedom. Slow drifts in the data were removed last, using a high pass filer (1/40 Hz cutoff). No additional spatial smoothing was applied to the data apart from the smoothing inherent to resampling and motion correction.

Signal amplitude time-series were normalized via Z-scoring on a voxel-by-voxel and run-by-run basis. Z-scored data were used for all further analyses unless mentioned otherwise. Trial events were jittered with respect to TR onsets, and unless mentioned otherwise, trial events were rounded to the nearest TR. To recover the univariate BOLD time courses for all three memory distractor conditions we estimated the Hemodynamic Response Function (HRF) for each voxel at each time point of interest (0–19.5 seconds from memory target onset). Estimated HRF’s were then averaged across all voxels in each ROI (see also **Supplementary Fig. 3**). Analyses performed after preprocessing was completed were all done in MATLAB 2016b using custom functions.

### Identifying Regions of Interest (ROIs)

To identify voxels that were visually responsive to the donut-stimuli, a General Linear Model (GLM) was performed on data from the mapping task using FSL FEAT (FMRI Expert Analysis Tool, version 6.00). Individual mapping runs were analyzed using BET brain extraction^30^ and data prewhitening using FILM^3^. Predicted BOLD responses were generated for blocks of “donut” and “circle” stimuli by convolving the stimulus sequence with a canonical gamma hemodynamic response function (phase = 0 s, sd = 3 s, lag = 6 s). The temporal derivative was also included as an additional regressor to accommodate slight temporal shifts in the waveform to yield better model fits and to increase explained variance. Individual runs were combined using a standard weighted fixed effects model. Voxels that were significantly more activated by the donut compared to the circle (p = 0.05; FDR corrected) were defined as visually responsive and used in all subsequent analyses.

Standard retinotopic mapping procedures^31,32^ were employed to define 9 a priori ROIs of early visual (V1–V3, V3AB, hV4) and parietal (IPS0–IPS3) areas. Retinotopic data were acquired during an independent scanning session, that utilized both meridian mapping techniques (with checkerboard “bowtie” stimuli shown alternating between the horizontal and vertical meridian) and polar angle techniques (with a slowly rotating checkerboard wedge) to identify the visual field preferences of voxels (stimuli described in more detail in^33^). Anatomical and functional retinotopy analyses were performed using a set of custom wrappers around existing FreeSurfer and FSL functionality. ROIs were combined across left and right hemispheres and across dorsal and ventral areas (for V2–V3) by concatenating voxels.

Only visually responsive voxels, selected using the localizer procedure described above, were included in the ROI of each retinotopic area. We only included data for retinotopic areas in which the number of visually responsive voxels exceeded 50 for every single participant.

### fMRI Analyses: Inverted Encoding Model

To reconstruct remembered and perceived orientations from voxel responses, an Inverted Encoding Model (IEM) was implemented^11,12^ with orientation as the feature dimension. The first step in this analysis is to estimate an encoding model using voxel responses in a cortical region of interest. These data are considered the “training set” (**Supplementary Fig. 4a**, left), and are combined with 9 idealized tuning functions, or “channels” (**Supplementary Fig. 4a**, right), to parameterize an orientation sensitivity profile for each voxel. The second step in the analysis combines the estimated sensitivity profiles in each voxel with a novel pattern of all voxel responses in a ROI on a single trial in the experimental data set (the “test set”, **Supplementary Fig. 4b**, left) to reconstruct the orientation that was remembered or viewed on that trial (**Supplementary Fig. 4b**, right). The IEM has the general form:

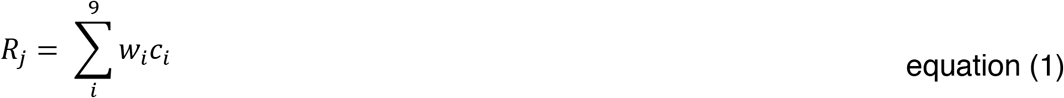

Where *R_j_* is the response *R* of voxel *j*, and *c_i_* is the channel magnitude *c* at the *i*^th^ of 9 channels. A voxel’s sensitivity profile over orientation space is captured by 9 weights *w*. Channels were modeled as:

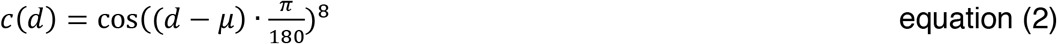

Where *d* is the distance in degrees from the channel center *μ*. Channel centers were spaced 20° apart.

For the first step of the IEM, equation 1 can be expressed as:

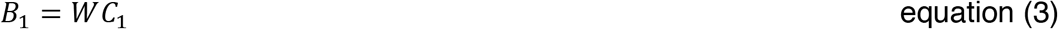

Here, a matrix of observed BOLD responses *B*_1_(*m* voxels x *n* trials) is related to a matrix of modeled channel responses *C_1_* (*k* channels x *n* trials) by a weight matrix *W* (*m* voxels x *k* channels). For each trial, *C_1_* is the pointwise product of a stimulus mask (i.e. “1” at the true stimulus orientation, “0” at all other orientations) with the idealized tuning functions. *W* quantifies the sensitivity of each voxel at each idealized orientation channel, and can be computed with least-squares linear regression:

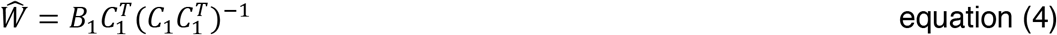

Estimating the sensitivity profiles concludes the first encoding step of the IEM. The second step of the IEM inverts the model, using the estimated sensitivity profiles of all voxels 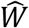 (*m* voxels x *k* channels) in combination with a “test set” of novel BOLD response data *B*_2_ (*m* voxels x *n* trials) to estimate the amount of orientation information at each channel 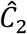 (*k* channels x *n* trials):

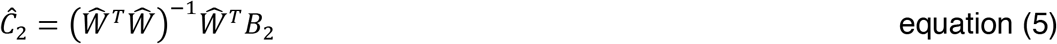

This step uses the Moore-Penrose pseudoinverse of 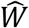, and it is multivariate in nature since it uses the sensitivity profiles across all voxels to jointly estimate channel responses 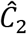 for each trial of the “test set”. This effectively forms a ‘reconstruction’ of the remembered or seen stimulus feature on a trial-by-trial basis.

Because grating orientations could take any integer value between 1° and 180°, both the encoding (equation 4) and inversion (equation 5) steps of the IEM were repeated 20 times. On each repeat, the centers of 9 idealized tuning functions were shifted by 1° (equation 2), and we estimated the channel responses 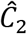 at those 9 centers, until the entire 180° orientation space was tiled. This procedure thus yielded estimated channel responses 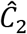 for each degree in orientation space.

Importantly, for all our IEM analyses we utilized independent data from the mapping task as the “training set” and data from the memory task as the “test set”. To obtain single trial activity estimates, mapping data were averaged across a time window of 4.8–9.6 seconds (6–12 TR’s) after donut onset, and the memory data over a window of 5.1–13.1 seconds (7–17 TRs) after target offset.

### Fidelity and behavioral metrics

Orientation reconstructions were quantified using a fidelity metric derived from basic geometry. At each degree in orientation space (wrapped onto a 2π circular space), we took the channel response, and projected this vector onto the center of the tuning function (i.e. at zero degrees) via 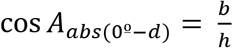, where *A* is the angle between the tuning function center (at 0°) and the degree in orientation space being evaluated (*d*), and *h* is the channel response at *d* (i.e. the hypotenuse of a right triangle). We thus project the length of the vector *h* onto *O°* by solving for *b* (i.e. the adjacent side of a right triangle). This procedure is repeated for all degrees in orientation space, after which we calculated the mean of all projected vectors. Thus, the projected vector mean reflects the amount of ‘energy’ at a remembered or sensed orientation (i.e. center of the reconstruction).

Behavioral data were fit using a ‘mixture model’^19,34^ which describes a distribution of response errors (**Supplementary Fig. 1a**) as a mixture of two components. The first is a von Mises distribution with concentration *κ* and center *μ*, describing responses under the assumption that a target item was in memory. The second is a uniform distribution represented by one parameter, *pU*, describing the proportion of responses under the assumption that a target item was not in memory. Thus, a change in *μ* would indicate any systematic shift of the distribution with respect to the correct response, a change in *κ* (which we report in units of *sd*) would indicate a change in memory precision, and a change in *pU* would indicate changes in the probability of memory failure. We calculated circular statistics using the circular statistics toolbox^35^.

### Statistical procedures

All statistical statements reported here were based on permutation testing over 1000 permutations. Note that this constrains the resolution of our p-values to a lower limit of p ≤ 0.001. To test if fidelity metrics were significantly greater than zero, we generated permuted null distributions of fidelities for each participant, ROI, and condition (and for each timepoint, in the analyses shown in **Fig. 1d** and **Supplementary Fig. 8**): On each permutation we first reshuffled target orientation labels for all trials before performing the inversion step of the IEM (effectively randomizing single trial reconstructions relative to the true orientation). Second, we calculated the fidelity for the trial-averaged orientation reconstruction in a manner identical the intact reconstructions. This resulted in one ‘null’ fidelity estimate per permutation. Combining six fidelities across participants resulted in one t-statistic per permutation 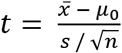, where 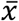 and *s* are the mean and standard deviation of amplitudes across participants, *μ*_0_ is the null hypothesis mean (i.e. 0), and *n* is the sample size. To test individual participant fidelities against zero (**Supplementary Fig. 6**) we directly compared the fidelity of each intact reconstruction against the permuted null distribution of fidelities for that subject, condition, and ROI. To test across-participant fidelities against zero (**Fig. 1c, 1d** and **Supplementary Fig. 8**) we compared the *t*-statistic calculated from the intact data against the permuted null distribution of t-statistics for that condition, ROI, and timepoint. Reported tests against zero were one-sided and uncorrected.

To test if there were differences in fidelity between ROI’s and conditions (**Fig. 1c**), we used within-subjects repeated-measures two-way ANOVA’s. First, we calculated the F-statistic from the intact data for the two main effects of condition and ROI, as well as their interaction. Next, we generated permuted null distributions of F by shuffling the condition and ROI labels per subject, and calculating the F-statistics for the main effects and the interaction on each permutation. Each data-derived F was then compared to its null distribution of F’s to get the p-value. Significant main effects were followed up with post-hoc paired-sample t-tests: We calculated the mean at each level of the factor showing the main effect (i.e. across the other factor) and performed pairwise comparisons between each of the factors, comparing the data derived 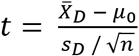, (with 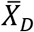 and *s_D_* denoting the mean and *sd* of the pairwise differences) against a permuted *t* distribution generated by reshuffling condition labels on each iteration. Significant interactions were followed up by first applying this randomization procedure to within-subject one-way ANOVA’s in each ROI, followed by similar randomization-based paired-sample t-tests. If this procedure did not reveal the source of the interaction, we compared each two ROI’s, looking for an interaction between ROI and condition using randomization and within-subject two-way ANOVA’s. All post-hoc interaction tests were performed as already described above.

### Glitches

S01 completed 4 sessions of scanning, but on the first day the projector settings had been changed such that we were presenting stimuli as ovals rather than circles. Data from this session has been excluded from analysis. S02 was also scanned 4 times, but data transfer after one of the sessions failed, and the data were deleted off the scanner center’s servers without having been backed up. For S04 we only collected 4 mapping runs on the first day of scanning because the scanner computer’s hard drive was full by the time we approached the end of the scan. The computer refused to do anything after that. On the first day of scanning S05 the scanner computer started deleting data files half way through. Consequently, data from the 2^nd^ mapping run was deleted and the 5^th^ memory run was aborted (with imaging data collection incomplete, while behavioral data collection was complete). To ensure exact counterbalancing in our analyses, an exact replica of this 5^th^ memory run was repeated as the first run on the second day of scanning.

## Supplementary Figures

**Supplementary Figure 1.**
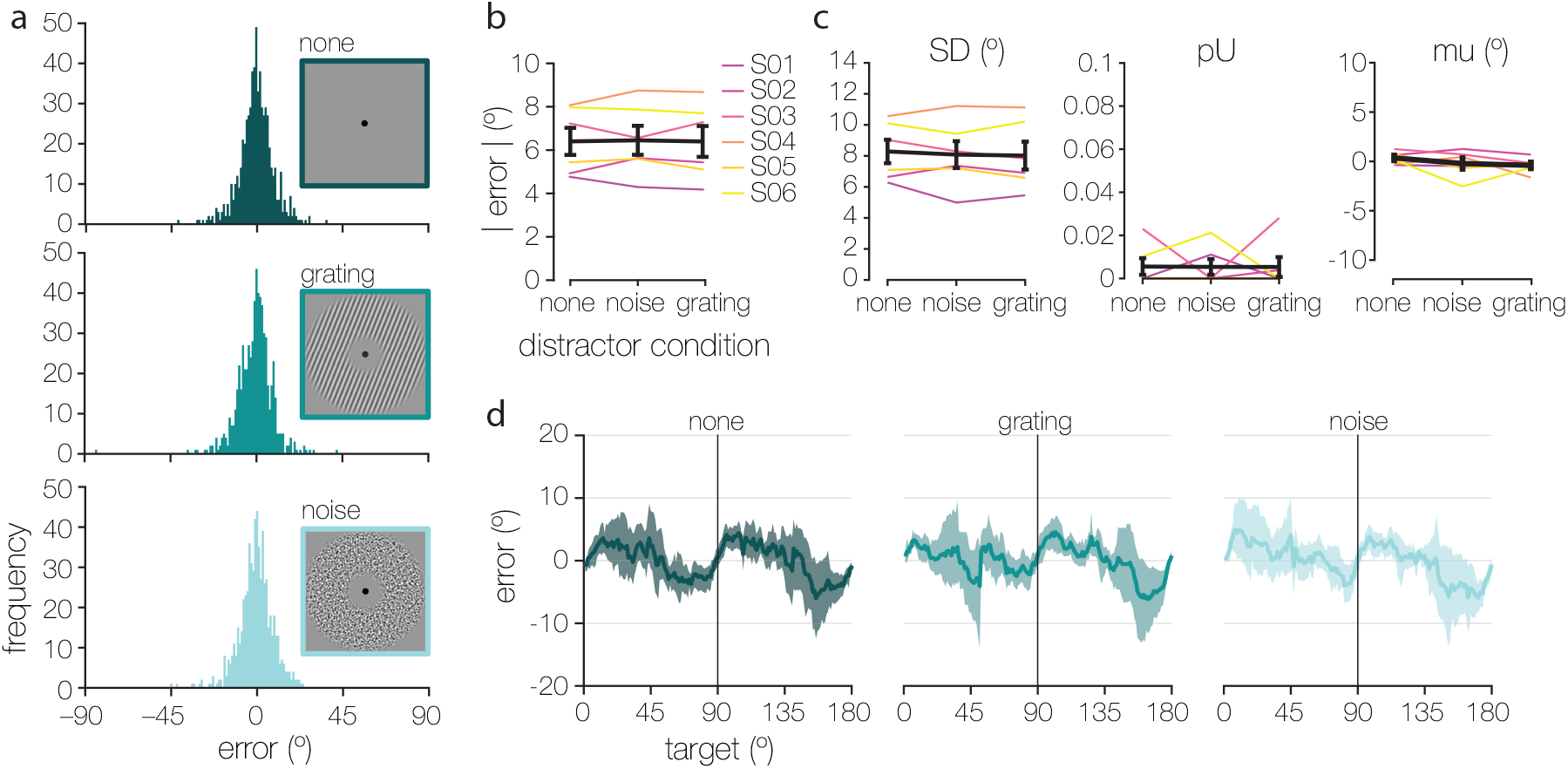
Behavioral performance during our working memory task in the scanner. (**a**) Distributions of behavioral error in degree, calculated by subtracting participant’s response from the target orientation on every trial. Data are sorted by distractor condition, with trials during which a Fourier filtered noise stimulus was shown during the delay in the top panel (lightest teal), trials during which a grating distractor was shown in the middle panel (mid teal), and trials with no distractor in the bottom panel (darkest teal) (**b**) There were no differences in the absolute behavioral error across the three distractor conditions, as indicated by a non-parametric one-way repeated-measures within-subject ANOVA (F_(2,10)_ = 0.044, p = 0.938). (**c**) When quantifying the error distributions in (a) with a commonly used ‘mixture model’^19^, no differences were observed in mnemonic precision as indexed by the mixture model standard deviation (leftmost panel; F_(2,10)_ = 0.476, p = 0.642), nor in the probability of memory failure as indexed by the proportion of uniform responses (middle panel; F_(2,10)_ = 0.001, p = 0.997). We furthermore found no differences in the circular mean (right-most panel; F_(2,10)_ = 1.399, p = 0.296). Error bars represent + 1 SEM. (**d**) Within-subject mean signed errors (on the y-axis) across orientation space (on the x-axis) for the three distractor conditions (in panels). A characteristic “bias away from cardinal”^20^ can be observed irrespective of distractor condition – shaded error areas represent bootstrapped 95% confidence intervals on the mean signed errors. The mean signed error at each degree was calculated within a window of ± 5±, thus smoothing the data within that range.

**Supplementary Figure 2.**
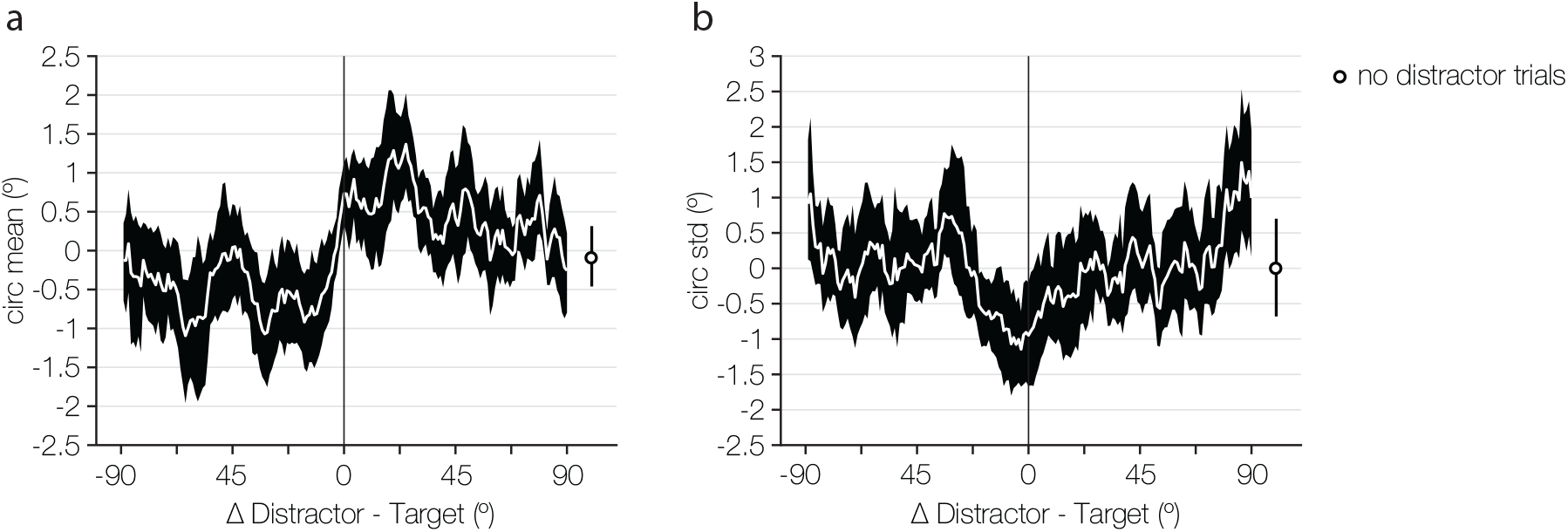
A psychophysical experiment of our distractor grating condition shows better behavioral performance when a target and distractor are similar versus dissimilar. Participants (N = 12; mean age = 19.33; 10 female) remembered a random target orientation, and an irrelevant distractor orientation was presented during the delay on 90% of trials. Target and distractor orientations were chosen randomly and independently as in the main experiment. However, in this experiment trials were faster (200ms target, 3000ms delay, 200ms distractor during the central portion of the delay, 800-1000ms jittered inter-trial interval), the response was unspeeded, and stimuli were smaller (2° radius, 2 c/°, 20% contrast, phase jittered). Participants performed a total of 1620 trials over the course of several days. (**a**) The mean response to the target as a function of the target-distractor similarity. Behavioral responses were attracted towards the irrelevant distractor orientation, and this attraction was most pronounced at target-distractor differences of around 22°. At its strongest, the attraction had a total magnitude of ~2°. The white line shows the circular mean at each target-distractor difference within a window of + 5°. Black error areas represent bootstrapped 95% confidence intervals on the within-subject circular means (across all possible target-distractor difference). The data point on the far right is the within-subject mean response when no distractor was presented during the delay. (**b**) The variability of participant’s responses covaried as a function of target-distractor similarity, in that memory was more precise for more similar orientations, and less precise for less similar orientations. The total magnitude of this effect was ~2°. The white line shows the circular standard deviation calculated within a ±5° window of the target-distractor difference. Black error areas are again 95% bootstrapped confidence intervals, and the data point on the far right the trials without distractors. Both the findings in (**a**) and (**b**) replicate previous work with irrelevant distractor orientations^13^, with the most notable difference in paradigms that here we had entirely independent target and distractor orientations, whereas in previous experiments there were dependencies between the two. While these effects are small, they far exceed the JND for orientation, and provide evidence for an interaction between the memory target and irrelevant distractors.

**Supplementary Figure 3.**
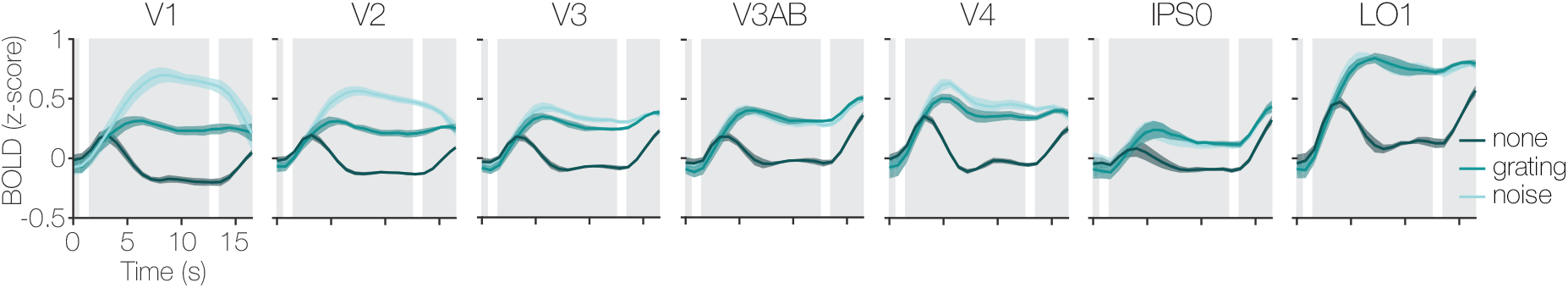
Hemodynamic response functions across all retinotopically defined ROIs (left-to-right panels) during the three distractor conditions (different shades of teal). Distractors presented during the delay effectively drove univariate response in all ROIs, with the Fourier filtered noise distractor (light teal) yielding especially strong activation in V1 and V2. The three gray panels represent the target, distractor, and recall epochs of the working memory trial (from left-to-right). Shaded error areas represent + 1 within-subject SEM.

**Supplementary Figure 4.**
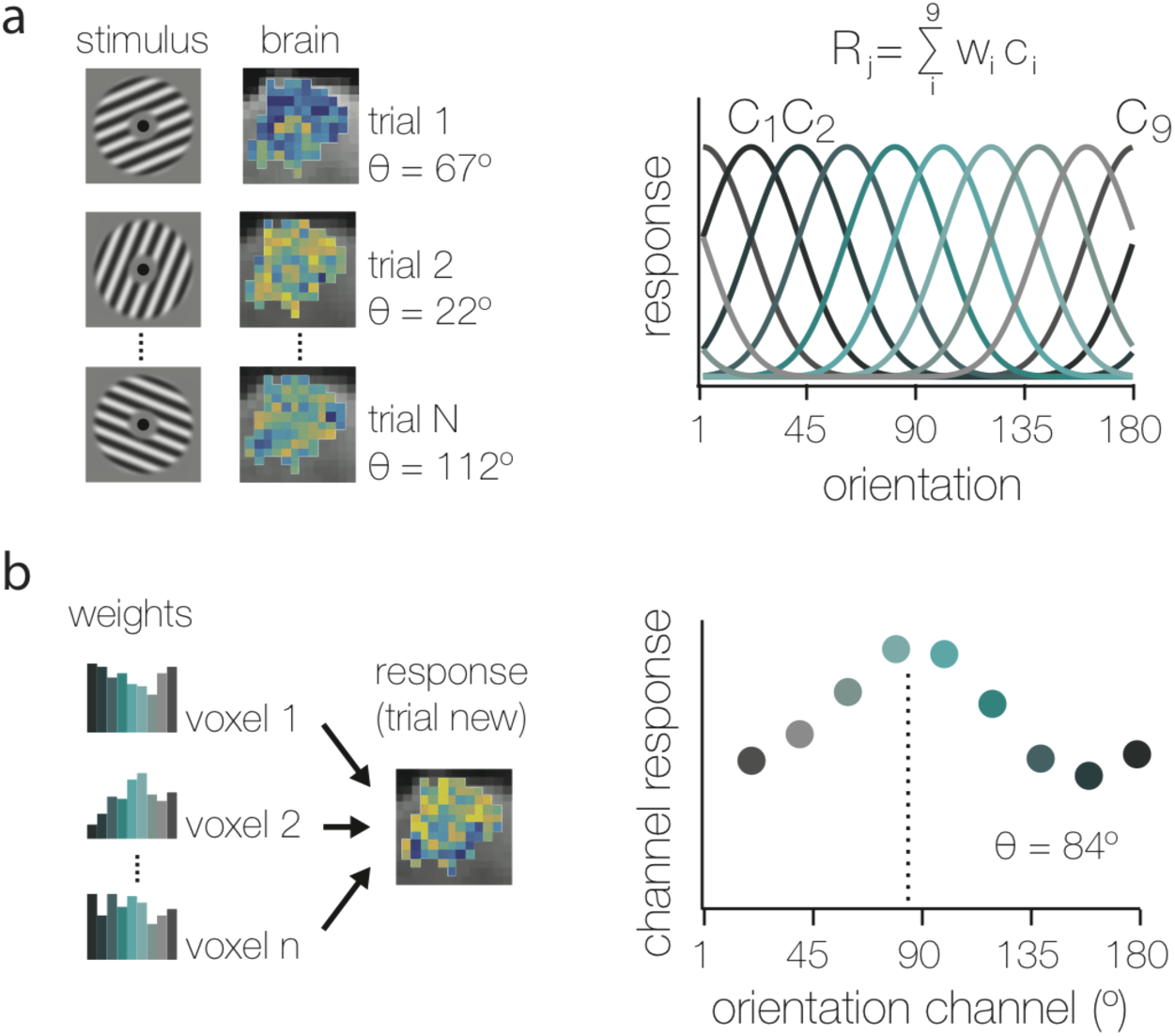
Inverse Encoding Model (IEM). (**a**) Estimating the encoding model is the first step in the IEM. Each voxel differs with respect to the orientation it preferentially responds to, and showing many orientations over many trials is used to quantify this response profile (left). Response profiles (R) for each voxel (j) are the weighted (w) sum of 9 hypothetical orientation channels (i), as shown on the right. (**b**) Inverting the encoding model is the second step in the IEM, and generates orientation reconstructions from voxel activity patterns. Channel weights represent each voxel’s orientation selectivity, and when a new response is evoked (left), the combined selectivity of all voxels is used to reconstruct this new orientation from the voxel pattern.

**Supplementary Figure 5.**
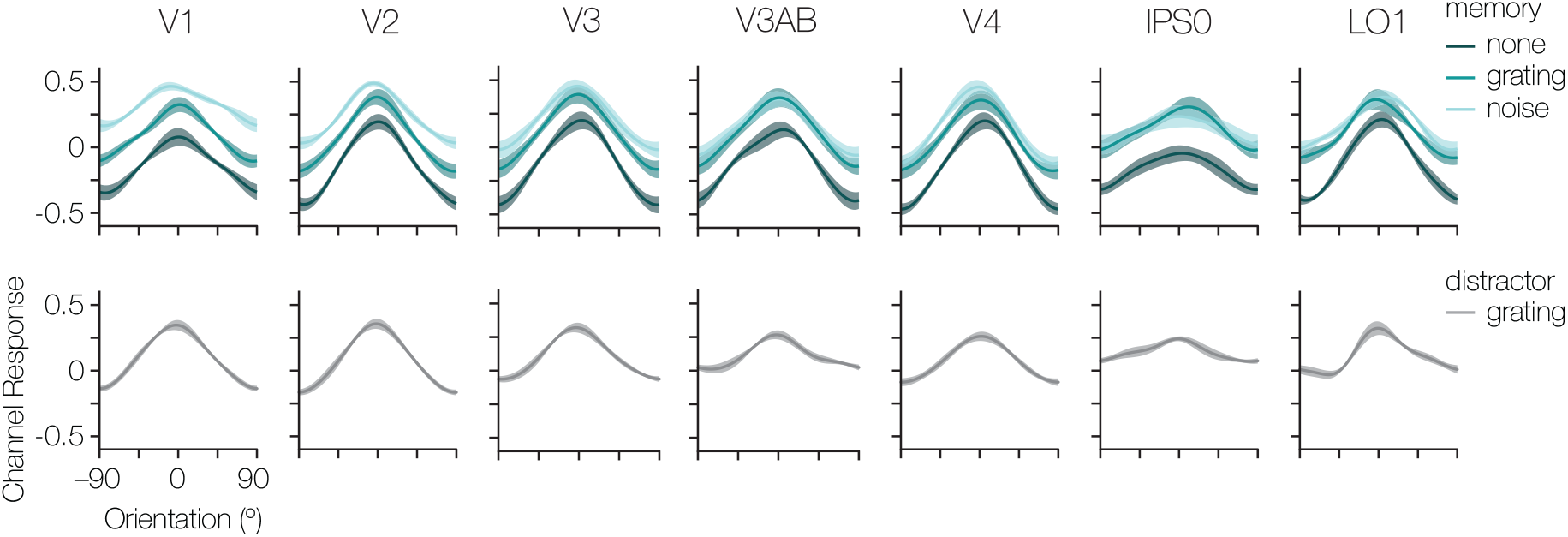
Reconstructions in all ROIs during the working memory delay (5.6–13.6 seconds after stimulus onset) for the orientation held in memory (top panels), and for the physically-present orientation on grating distractor trials (bottom panels). Reconstructions for V1 are also shown in Fig. 1b but are included here as well for ease of comparison. Shaded error areas represent ± 1 within-subject SEM.

**Supplementary Figure 6.**
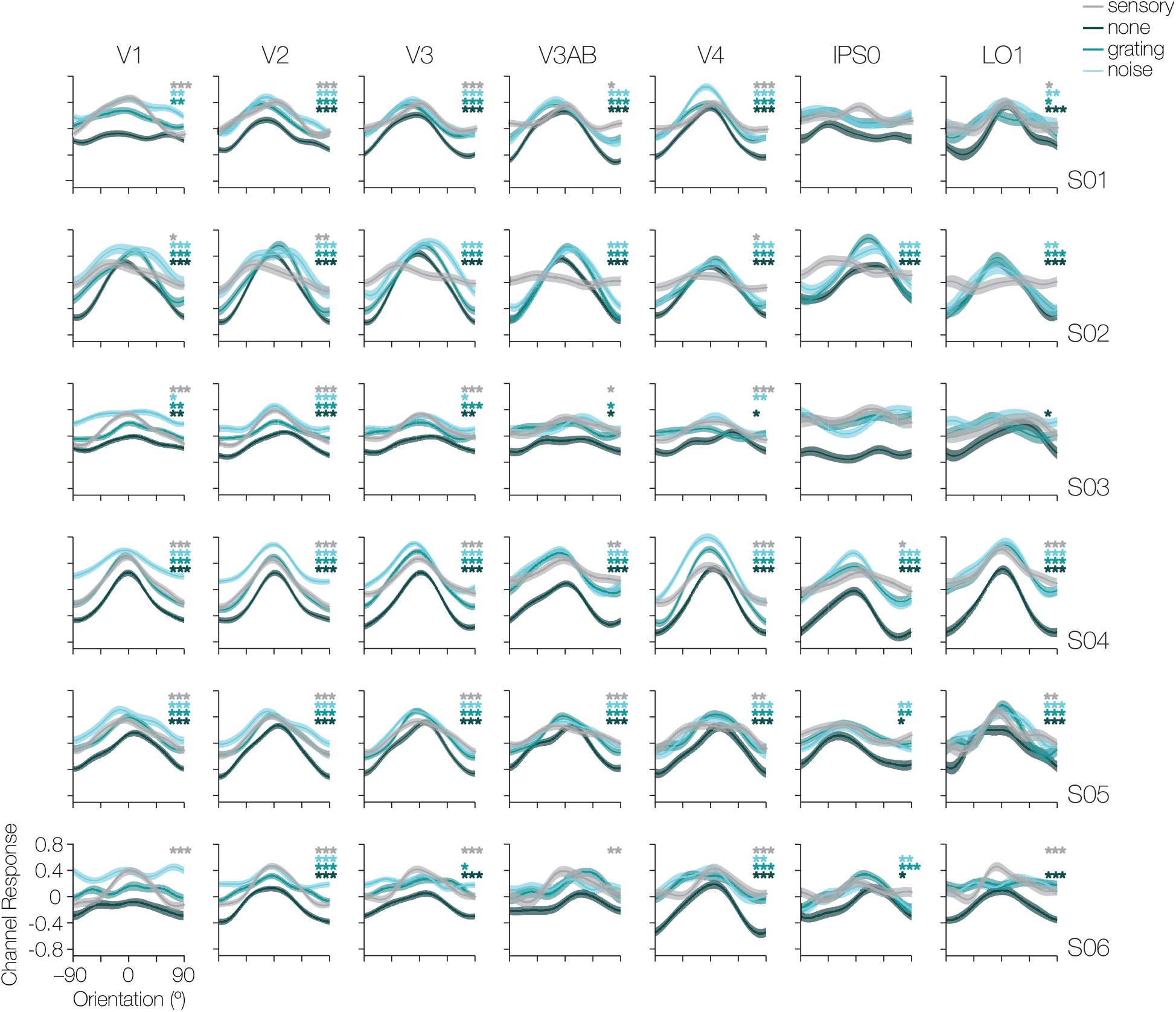
Reconstructions as in Supplementary Fig. 5 shown for all individual participants, with remembered orientations in teal hues, and physically-present sensory distractor orientations in grey. Asterisks in each subplot panel indicate significance of the reconstructed fidelity (indexed by a fidelity > 0) for that participant and ROI. Asterisk color indicates the condition referred to, and asterisk quantity indicates the level of significance with one, two, and three asterisks representing p-values of p ≤ 0.05, p ≤ 0.01, and p ≤ 0.001, respectively. Shaded error areas represent ± 1 SEM.

**Supplementary Figure 7.**
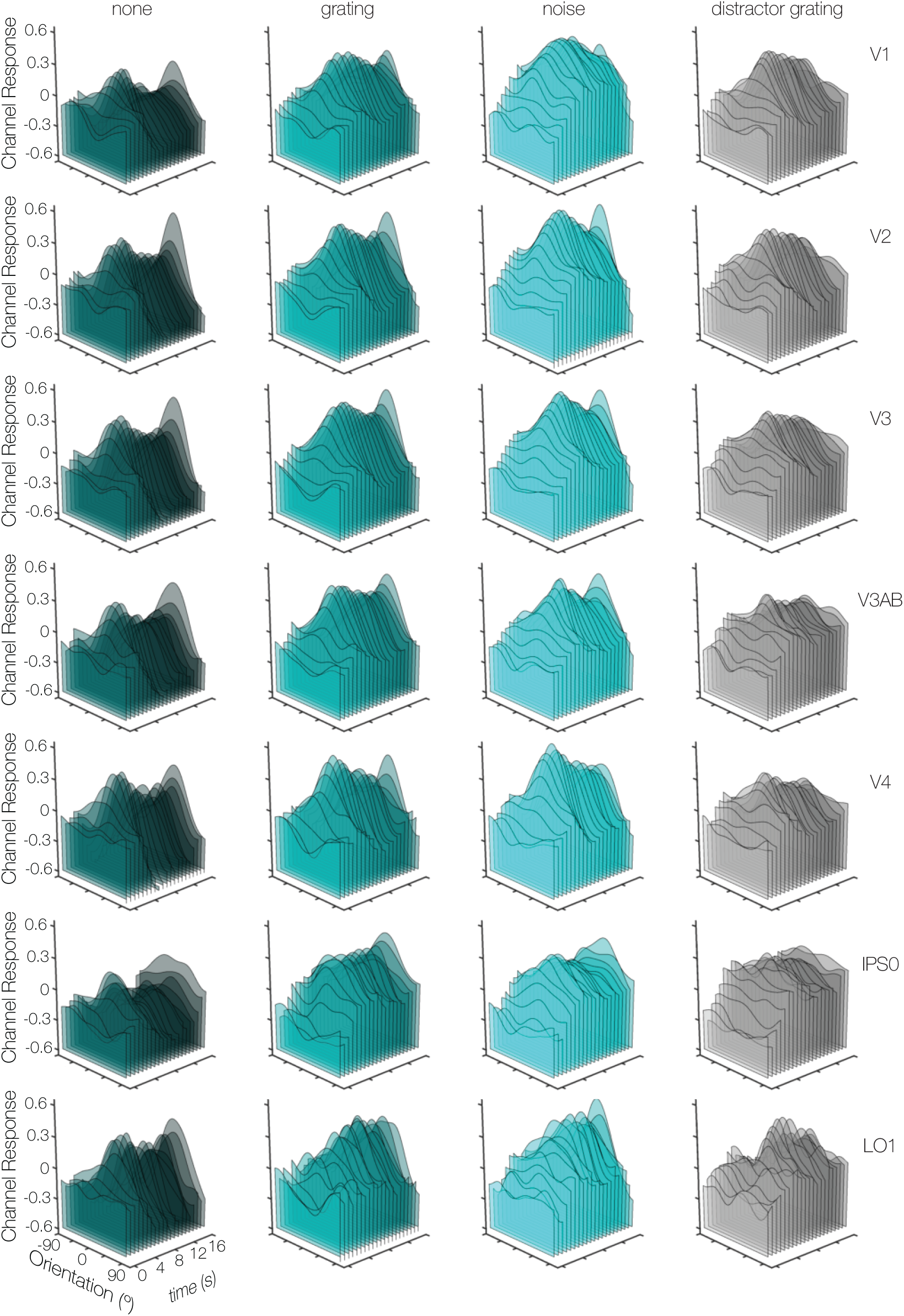
Reconstructions of remembered orientations and distractor orientations over time. The time axis starts at “0” which is trial onset, and shows the mean reconstruction across participants at each 800ms TR (for a total of 21 TR’s, including time “0”). Reconstructions for the remembered orientations are shown for each distractor condition in the first 3 rows (shades of teal), and reconstructions for the physically-present distractor orientation is shown in the last row (grey).

**Supplementary Figure 8.**
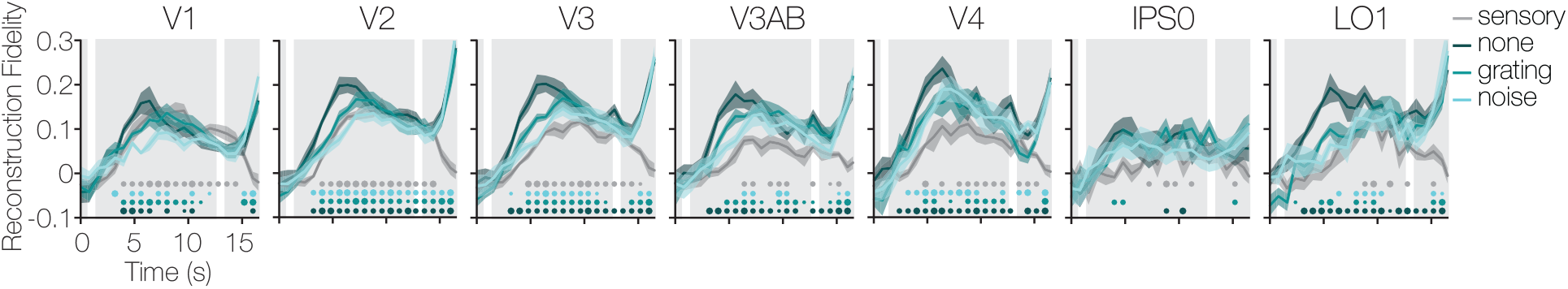
Memory fidelity was calculated for orientation reconstructions at each timepoint during a trial, plotted here in steps of 0.8s TR’s along the x-axis. The three gray panels represent the target, distractor, and recall epochs of the working memory trial (left-to-right). Dots in the bottom of each plot indicate the time points at which the fidelity was significantly above zero. Small, medium, and large dot sizes indicate significance levels of p ≤ 0.05, p ≤ 0.01, and p ≤ 0.001, respectively. A significant fidelity for the remembered orientation arises about 3–4 seconds into the trial irrespective of distractor condition (in shades of teal), and persists throughout the delay. On trials with a grating distractor, the physically-present orientation is also represented throughout (though arising a little later, around 4–5 seconds into the delay, consistent with its delayed onset of 1.5 seconds after the target), and dissipates roughly 2 seconds after its offset. Fidelity over time for V1 was also shown in **Fig. 1d** and is included here for ease of comparison. Shaded error areas represent ± 1 within-subject SEM.

**Supplementary Figure 9.**
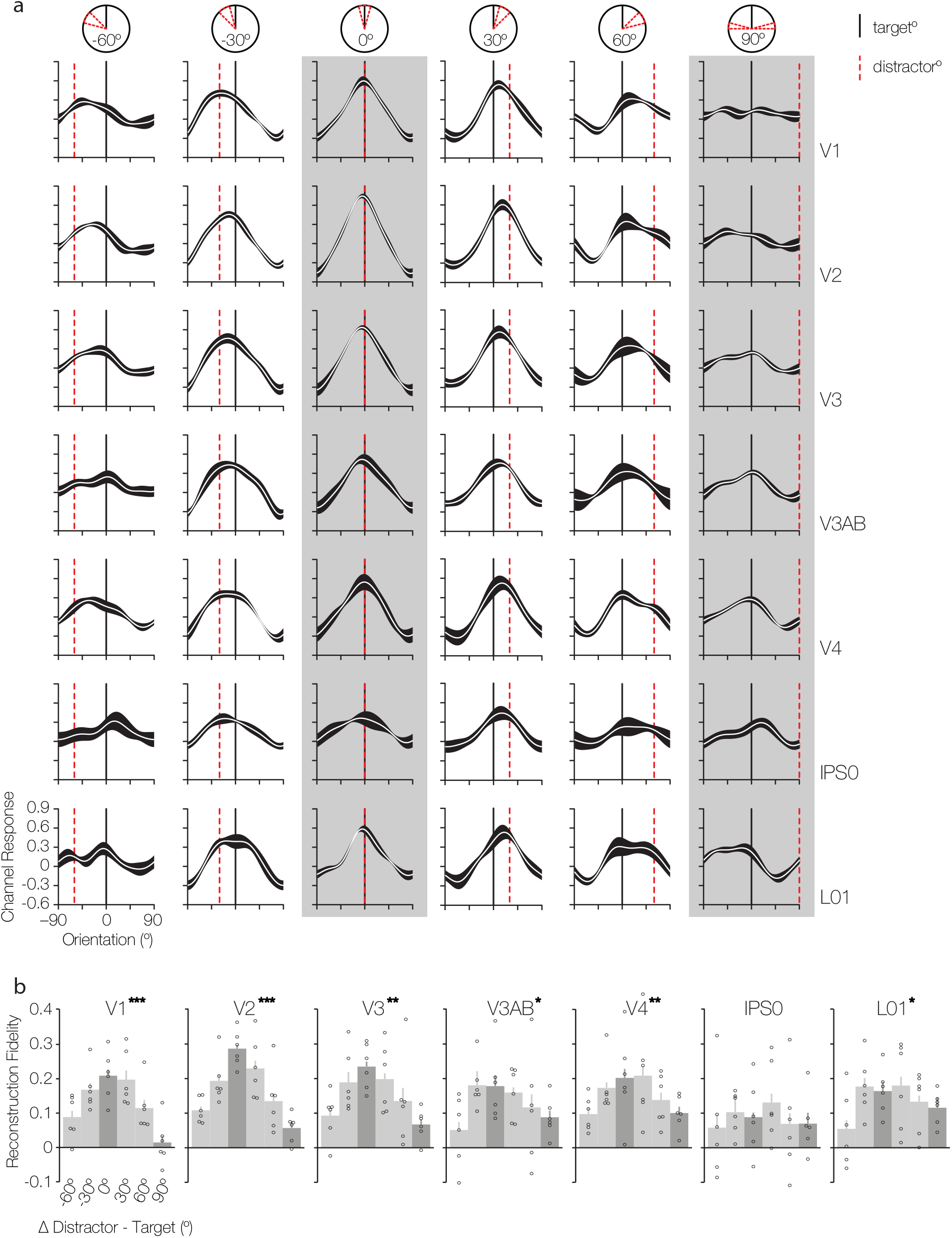
Increased reconstruction fidelity when target and distractor are similar versus dissimilar. (**a**) Data in the grating distractor condition were binned according to the target-distractor orientation similarly (in columns) for each ROI (in rows). Bins centers were chosen in 30° steps between maximal (0°) and minimal (90°) target-distractor similarity (columns depicting minimum and maximum similarity are highlighted my gray panels). The black and dashed-red vertical lines represent the memory target orientation, and bin-center of the distractor orientation, respectively. For every participant, all trials within a bin were averaged and the fidelity (see **b**) was determined by projecting all vectors onto the circular mean across subjects. These are the group averaged reconstructions (black error areas indicate + 1 within-subject SEM). (**b**) Fidelity of the orientation reconstructions. Following up on a significant interaction between similarity-bin and ROI (F_(30,150)_ = 1.802, p = 0.011), one-way ANOVA’s in each ROI showed that especially at the earliest levels of visual processing (V1–V3) there are interdependencies between target and distractor (one, two, or three asterisks indicate significance in each ROI of p ≤ 0.05, p ≤ 0.01, and p ≤ 0.001, respectively). These interdependencies make it hard to separately estimate information from the target and the distractor as they co-vary, and the superimposed reconstructions cancel each other out. Nevertheless, this analysis shows that fidelity is enhanced when target and distractor are more similar, and highlights local interactions between memory and sensory information in early visual cortex.

## Supplementary Tables

**Supplementary Table 1.** Post-hoc comparisons of the main effect of distractor condition on fidelity. Each cell holds the t- and p-value for the comparison between two conditions (collapsed across ROI). White cells indicate a significant difference between conditions.

|  | none | grating | noise |
| --- | --- | --- | --- |
| none |  |  |  |
| grating | t = 2.164<br>p = 0.074 |  |  |
| noise | t = 6.298<br>p < 0.001 | t = 3.398<br>p = 0.01 |  |

**Supplementary Table 2.** Post-hoc comparisons of the main effect of ROI on fidelity. Each cell holds the t- and p-value for the comparison between two ROIs (collapsed across distractor condition). White cells indicate a significant difference in ROI.

|  | V1 | V2 | V3 | V3AB | V4 | IPS0 | L01 |
| --- | --- | --- | --- | --- | --- | --- | --- |
| V1 |  |  |  |  |  |  |  |
| V2 | t = 5.767<br>p < 0.001 |  |  |  |  |  |  |
| V3 | t = 4.159<br>p < 0.001 | t = 0.305<br>p = 0.772 |  |  |  |  |  |
| V3AB | t = 1.731<br>p = 0.108 | t = 0.718<br>p = 0.442 | t = 1.136<br>p = 0.27 |  |  |  |  |
| V4 | t = 2.637<br>p = 0.03 | t = 0.772<br>p = 0.376 | t = 0.574<br>p = 0.478 | t = 1.053<br>p = 0.252 |  |  |  |
| IPS0 | t = 0.898<br>p = 0.366 | t = 2.901<br>p < 0.001 | t = 2.617<br>p = 0.07 | t = 1.7<br>p = 0.126 | t = 3.05<br>p < 0.001 |  |  |
| L01 | t = 3.143<br>p < 0.001 | t = 1.281<br>p = 0.272 | t = 2.196<br>p = 0.06 | t = 0.271<br>p = 0.724 | t = 1.58<br>p = 0.184 | t = 1.751<br>p = 0.13 |  |

**Supplementary Table 3.** First pass post-hoc comparisons of the interaction effect between sensory condition (i.e. remembered or physically-present grating) and ROI on fidelity. Each cell holds the t- and p-value for the comparison between remembered and physically-present amplitudes for each ROIs. No significant differences were found in any of the ROIs.

| V1 | V2 | V3 | V3AB | V4 | IPS0 | L01 |
| --- | --- | --- | --- | --- | --- | --- |
| t = 0.508<br>p = 0.572 | t = 0.256<br>p = 0.806 | t = 1.139<br>p = 0.308 | t = 1.635<br>p = 0.106 | t = 1.736<br>p = 0.148 | t = 0.883<br>p = 0.38 | t = 0.677<br>p = 0.558 |

## Notes

**Conflict of interest**: The authors declare no conflict of interest

